# Ythdf2 regulates cardiac remodeling through its m^6^A-mRNA target transcripts

**DOI:** 10.1101/2022.12.16.520765

**Authors:** V. Kmietczyk, J. Oelschläger, E. Varma, S. Hartl, M. Konstandin, A. Marx, P. Gupta, Z. Loewenthal, V. Kamuf-Schenk, L. Jürgensen, C. Stroh, A. Gorska, T. Jakobi, N. Frey, M. Völkers

## Abstract

m^6^A mRNA methylation controls cardiomyocyte function and increased overall m^6^A levels are a stereotyping finding in heart failure independent of the underlying etiology. However, it is largely unknown how the information is read by m^6^A reader proteins in heart failure. Here we show that the m^6^A reader protein Ythdf2 controls cardiac function and identified a novel mechanism how reader proteins control gene expression and cardiac function. Deletion of Ythdf2 in cardiomyocytes *in vivo* leads to cardiac hypertrophy, reduced heart function, and increased fibrosis during pressure overload as well as during aging. Similarly, *in vitro* the knockdown of Ythdf2 results in cardiomyocyte growth and remodeling. Mechanistically, we identified the eucaryotic elongation factor 2 as a major target of Ythdf2 using cell type specific Ribo-seq data. Our study expands our understanding on the regulatory functions of m^6^A methylation in cardiomyocytes and how cardiac function is controlled by the m^6^A reader protein Ythdf2.

## Introduction

N^6^-methyladenosine (m^6^A) mRNA modification controls diverse cellular processes in mammalian cells by affecting all aspects of the mRNA’s fate [1]. In the heart, levels of m^6^A are comparably low in comparison to liver, kidney and brain [2]. However, in human failing hearts as well as various animal models of heart failure overall levels of m^6^A increase compared to non-failing myocardium [3–6]. Studies using knockout and overexpression of the N6-adenosine-methyltransferase 70 kDa subunit (Mettl3) mRNA or the m^6^A eraser fat mass and obesity-associated protein (FTO) have proven that m^6^A represents an additional layer of gene expression control on the transcript level in the heart which controls cardiac growth and cardiomyocyte contractility [3–5, 7]. Those studies clearly showed the impact of mRNA modifications on cardiac function. However, how the information of methylated transcripts is transformed by m^6^A-binding proteins (‘m^6^A readers’), and how the reader proteins affect cardiac function is largely unknown.

Different protein families have been shown to function as m^6^A readers. The best studied protein family are the YT521-B homology (YTH) domain containing proteins-Ythdf1, 2 and 3 [8]. Ythdf1 was shown to boost translation without affecting the mRNA abundance [9], Ythdf3 is known to have similar effects as Ythdf1 and synergizes with it [10, 11], while Ythdf2 mainly induces decay thus decreasing the transcripts stability [12–14]. Various studies support the hypothesis that different species as well as different tissues and cell types choose different m^6^A reader proteins to exert specific m^6^A reading functions [15, 16]. Opposing to this hypothesis, Zaccara et al. suggest that all Ythdf proteins share their mRNA targets and work together in regulating the decay of the transcripts [17].

Very little is known about the m^6^A readers in the heart. Zhang et al. showed that the expression of m^6^A reader Ythdf2 is increased in hearts of heart failure patients [18]. Similarly, Xu et al. report an increase of Ythdf2 in human heart failure samples and genetic overexpression of Ythdf2 in mice protected against heart failure in response to transaortic constriction surgery (TAC) model [19], but a characterization of a cardiomyocyte-specific Ythdf2 KO mice is missing.

Therefore, the aim of this study was to uncover the role of Ythdf2 in cardiac remodeling, development of heart failure and unravel the underlying mechanism how reader proteins regulate cardiac gene expression and function. We found that the loss of Ythdf2 in cardiomyocytes induces cardiac remodeling *in vitro* and *in vivo*. Mechanistically, we identified by cell-type specific Ribosomal Profiling (Ribo-Seq) transcripts that are controlled by Ythdf2. Among them, the eucaryotic elongation factor 2 mRNA (*Eef2*) binds to Ythdf2 and Ythdf2 suppresses translation of Eef2. In line, Eef2 expression is increased upon loss of Ythdf2 by translational regulation. Our present study confirms the regulatory function of m^6^A methylation on cardiac function and identified a mechanism how the reader protein Ythdf2 promotes cardiac growth and remodeling.

## Results

### Ythdf2 regulates cell growth in primary cardiomyocytes

Previous studies using gain of/loss of function models of the methyltransferase Mettl3 or the de-methylase FTO have shown that m^6^A affects cardiac function and remodeling, but m^6^A reader proteins have not been comprehensively characterized in cardiomyocytes. Hence, we aimed to study the role of Ythdf2 which has been shown to regulate mRNA stability and translation [12, 20]. First, we aimed to study the effect of Ythdf2 depletion *in vitro* on protein synthesis and cell growth of neonatal rat ventricular myocytes (NRVMs), due to a positive correlation between protein synthesis and hypertrophy that has been established [21, 22]. We depleted Ythdf2 using small interfering RNA (siRNA) in NRVMs and induced hypertrophy by treatment with the a-adrenergic agonist phenylephrine (PE).

Successful depletion of Ythdf2 in NRVMs was validated by immunoblots **(Fig. 1A)** and a puromycin incorporation assay was performed to evaluate the translational activity of the cardiomyocytes. As expected, the treatment with PE increased puromycin incorporation indicating a higher protein synthesis rate. Ythdf2 depletion also significantly elevated the puromycin incorporation compared to controls **(Fig. 1 B)**.

**Figure 1.**
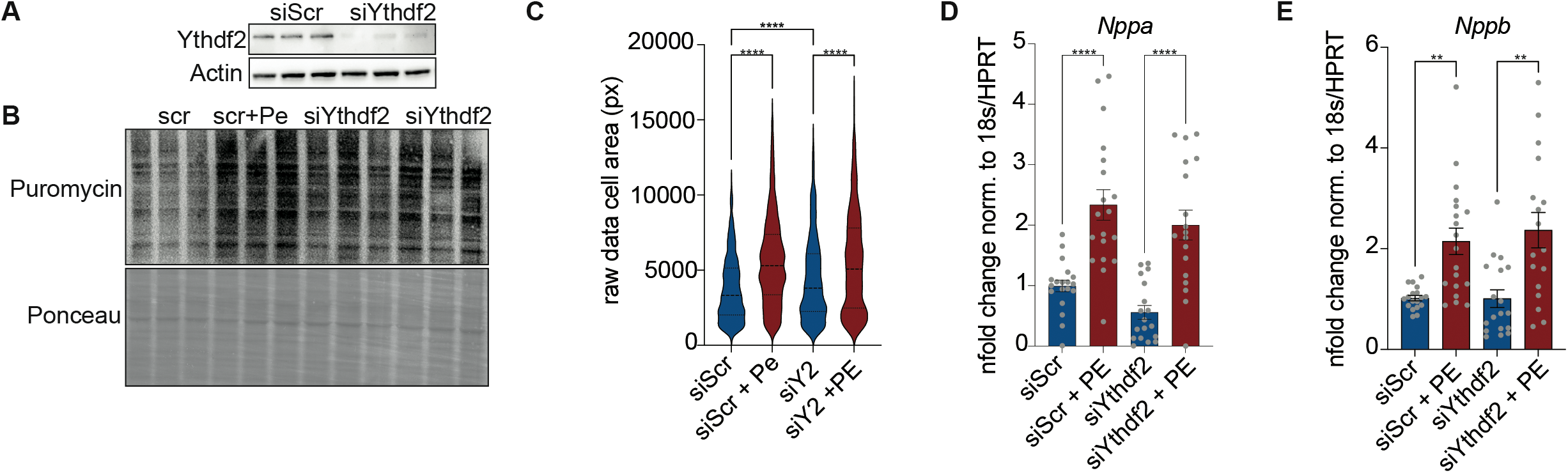
The knockdown of Ythdf2 in NRVMs increases cell size. **A** Western Blot showing the successful knockdown of NRVMs transfected with siRNA against Ythdf2. **B** Puromycin incorporation assay of NRVMs depleted of Ythdf2 and treated with PE, Ythdf2 KD increases translation. **C** Cell size quantification of the cell surface area, measured by In Cell Analyzer, ****P < 0.0001 by Kruskal Wallace test. mRNA expression levels of **D** *Nppa* and **E** *Nppb* in NRVMs for indicated groups, measured by RT-qPCR, ** P < 0.01, ****P < 0.0001 by one-way ANOVA. Scr-scrambled siRNA as control. Error bar indicates SEM

Next, cell growth was analyzed by immunostaining for desmin and measured using the automatic and high throughput IN Cell Analyzer imaging system with the C-MORE software [23]. PE induced cardiomyocyte growth compared to vehicle-treated cells as expected. Knockdown of Ythdf2 itself increased cell size significantly in comparison to the control cells transfected with scrambled (siScr) siRNA. Further stimulation with PE led to additional cell size increase to a similar extent compared to the control NRVMs **(Fig. 1 C)**.

Analysis of the RNA levels of the hypertrophy marker genes Atrial natriuretic peptide (*Nppa*) and brain natriuretic peptide (*Nppb*) by RT-qPCR showed a significant increase after PE treatment in control cells, but Ythdf2 depletion did not lead to any changes in expression of *Nppa* nor *Nppb* compared to control NRVMs treated with PE **(Fig. 1 D + E)**.

Since several studies showed that the depletion of one of the Ythdf paralogs leads to the upregulation of the other two Ythdf protein paralogs and moreover all three Ythdf proteins might share cellular same function and binding partners [17], the expression of the two other Ythdf protein paralogs, Ythdf1 and Ythdf3, was analyzed on protein level by western blotting and on the mRNA level by RT-qPCR. Ythdf1 and Ythdf3 protein as well as mRNA levels did not change significantly in the Ythdf2 depleted NRVMs in comparison the control siRNA transfected cells **(Suppl Fig. 1 A-D)**.

### Cardiomyocyte specific loss of Ythfd2 drives cardiac remodeling

As Ythdf2 depletion did affect primary cardiomyocyte growth *in vitro*, we continued to study the role of Ythdf2 during cardiac remodeling and hypertrophy *in vivo*. We obtained a floxed Ythdf2 mouse and crossed it with our Cre α-MHC/Ribotag mouse that expresses an HA-tag on the ribosomal protein RPL22 in cardiomyocytes to obtain a cardiomyocyte-specific Ythdf2 knockout, enabling isolation of cardiomyocyte-associated ribosomes for Ribo-seq (Suppl. Fig 2 A) [16]. For simplification the cardiomyocyte specific Ythdf2 knockout Ribotag mice used in this study are called Ythdf2 KO mice or KO mice in the following text.

We first controlled the expression of *Ythdf1* and *Ythdf3* in the Ythdf2 knockout mice by RT-qPCR. At an age of 14 weeks as well as with 30 weeks of age both transcripts are not significantly different expressed in KO mice compared to WT mice **(Fig. 2 A + B and E+F)**. We further examined the expression of the fetal gene markers *Nppa* and *Nppb* and Collagen, type I, alpha 1 (*Col1a1*) and Fibronectin (*FN1*) as fibrosis markers in the young adult (14 weeks) and old adult (30 weeks) mice by RT-qPCR. None of the transcripts were differentially expressed in 14 weeks old KO mice **(Fig. 2 C+D and Suppl. Fig. 2 B+C)**, however, in 30 weeks old mice hypertrophy and fibrosis markers were highly upregulated **(Fig. 2 G+H and Suppl. Fig. 2 D+E)**.

**Figure 2.**
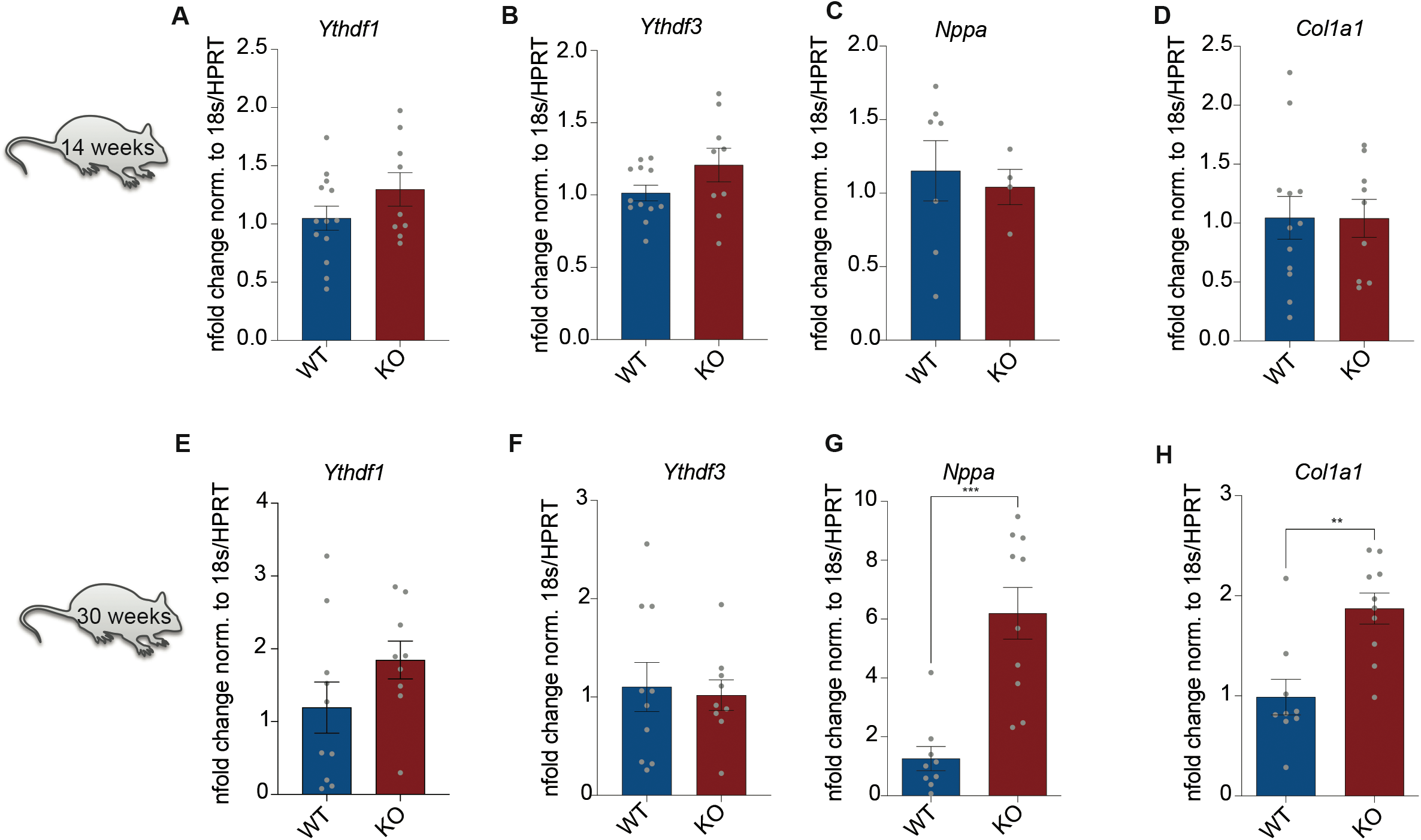
The cardiomyocyte-specific KO of Ythdf2 increases the expression cardiac remodeling markers in old mice. **A-D** mRNA expression levels of **A** *Ythdf1* **B** *Ythdf3* **C** *Nppa* and **D** *Col1a1* in left ventricles from 14 weeks old WT and Ythdf2 KO mice measured by RT-qPCR. **E-H** In 30 weeks old Ythdf2 KO mice, the mRNA expression levels of **E** *Ythdf1* **F** *Ythdf3* **G** *Nppa* and **H** *Col1a1* in comparison to WT mice at the same age, measured by RT-qPCR, ** P < 0.01, ***P < 0.0005 by unpaired t-test. Error bar indicates SEM

To further study the effect of Ythdf2 loss during cardiac remodeling we subjected WT and Ythdf2 KO mice to transverse aortic constriction (TAC) surgery [24]. In WT as well as KO mice ejection fraction decreased over time in response to TAC (**Fig. 3 A)**. Both, ejection fraction and fractional shortening, were significantly decreased in Ythdf2 KO mice compared to WT mice after 4 weeks after TAC surgery **(Fig. 3 A + B)**. Mice were sacrificed 4 weeks after surgery to assess heart dimensions, cardiomyocyte size, fibrosis, and gene expression. WT as well as KO mice displayed an increase in LV/body weight ratio, however there was no significant difference between these two groups **(Suppl. Fig 3 A)**. Immunostaining with wheat germ agglutinin (WGA) on heart sections from WT and KO mice after Sham or TAC surgery revealed an increase of cardiomyocytes cross-sectional areas (CSA) in WT after TAC surgery and an even further increase in Ythdf2 KO mice compared to WT TAC mice **(Fig. 3 C + D)**.

**Figure 3.**
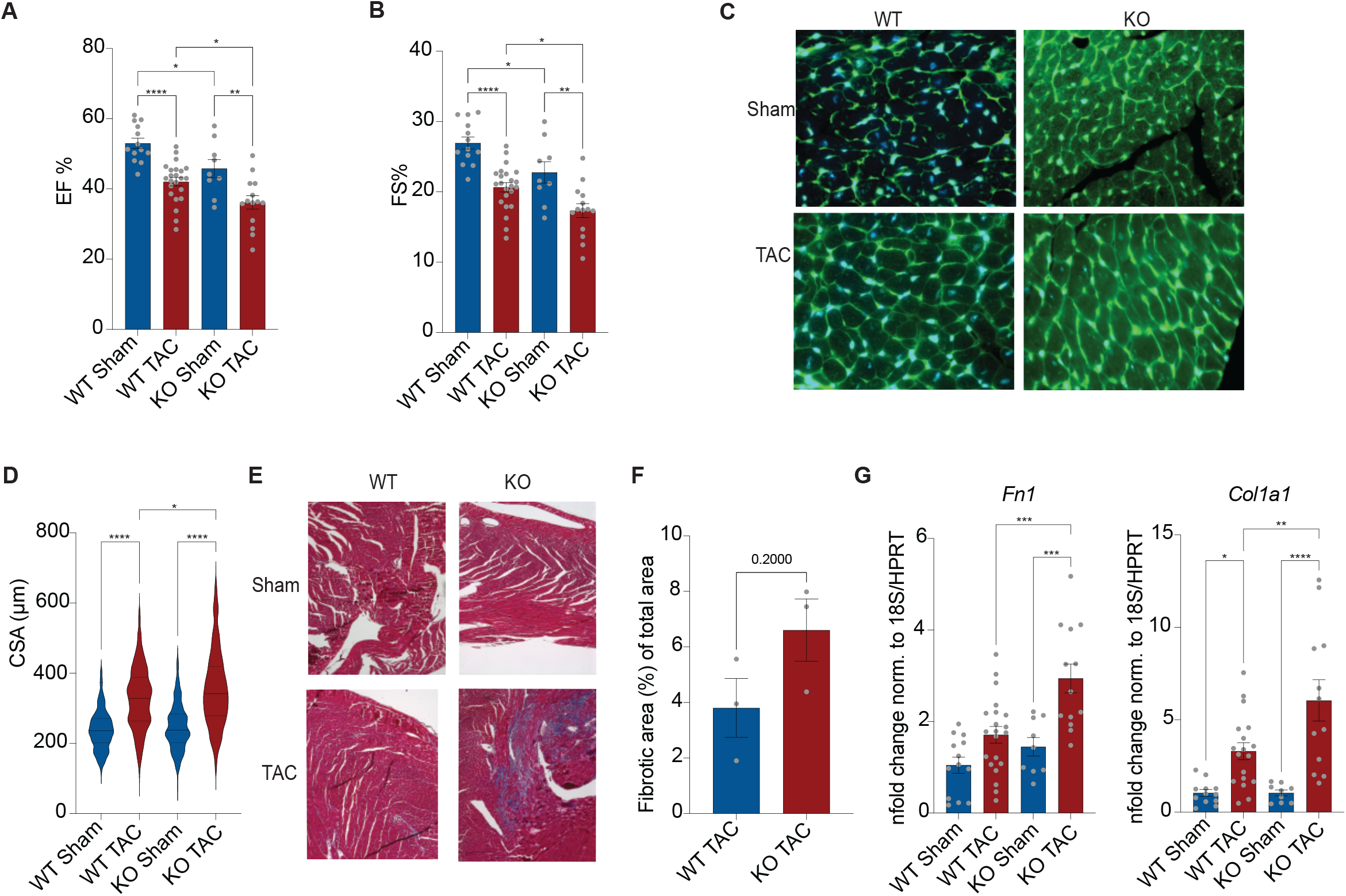
The cardiomyocyte-specific KO of Ythdf2 decreases heart function and increases signs of cardiac remodeling in young adult mice after TAC surgery. **A** Ejection fraction and **B** fractional shortening 4 weeks after TAC surgery in WT and in Ythdf2 KO mice measured by echocardiography, * P < 0.05, ** P < 0.01, **** P < 0.001 tested by one-way ANOVA. **C** Immunohistochemistry staining of heart sections from WT and KO mice after Sham or TAC surgery, stained for membrane proteoglycans by WGA (green) and for nuclei by DAPI (blue) to visualize the cross-sectional area of the cardiomyocytes. **D** Quantification of the cross-sectional cell surface area. * P < 0.05, **** P < 0.001 by Kruskal Wallace test. **E** Masson Trichrome staining of heart sections from WT and KO mice after Sham or TAC surgery. **F** Quantification of fibrotic areas by Image J, n = 3 mice, by unpaired t-test. **G** mRNA expression levels of the fibrotic markers *Fn1* and *Col1a1* in KO mice after TAC surgery in comparison to WT mice after TAC surgery, measured by RT-qPCR, * P < 0.05, ** P < 0.01, *** P < 0.005, **** P < 0.001 by one-way ANOVA. Error bar indicates SEM

Masson trichrome staining on heart sections showed the development of interstitial as well as patches of fibrosis in WT and KO mice after TAC surgery, and the percentage of fibrotic areas was higher in Ythdf2 KO mice in comparison to WT mice **(Fig. 3 E + F)**. Additionally, quantification by RT-qPCR showed that *Col1a1* and *Fn1*, as indicators for fibrosis, were significantly increased in Ythdf2 KO mice compared to WT mice **(Fig. 3 G)**.

Finally, hypertrophy marker gene expression was analyzed by RT-qPCR in WT and KO mice after TAC surgery. *Nppa, Nppb*, and myosin heavy chain 7 (*Myh7*) were elevated after 4 weeks TAC surgery in WT and KO mice, with no statistical difference between the experimental groups **(Suppl. 3 B-D)**.

In summary, Ythdf2 reduction in primary cardiomyocytes leads to cardiomyocyte growth. *In vivo*, Ythdf2 deletion leads to reduced heart function in aged mice and a gene expression profile indicating cardiac remodeling. Pressure overload in response to TAC surgery resulted in deteriorated heart function associated with increased fibrosis compared to WT mice.

### Identification of Ythdf2 targets

Our data suggest that Ythdf2 regulates cardiac remodeling by affecting cardiomyocyte growth and function. The next aim of this study was to identify mRNA targets of Ythdf2 that play a role in these processes. Hence, we performed Ribo-seq, from isolated ribosomes from left ventricles of our Ythdf2 KO mice and WT mice. WT and KO mice were crossed with the Ribo-tag mouse that express a HA-tagged RPL22 in cardiomyocyte, which enables the isolation of ribosomes from cardiomyocytes (**Fig. 4 A-B)**. We kept only periodic fragment lengths that showed a distinctive triplet periodicity for downstream analysis [25]. The obtained libraries yielded around 5 million usable periodic reads **(Suppl. Fig 4)**, showing high correlation between the replicates.

**Figure 4.**
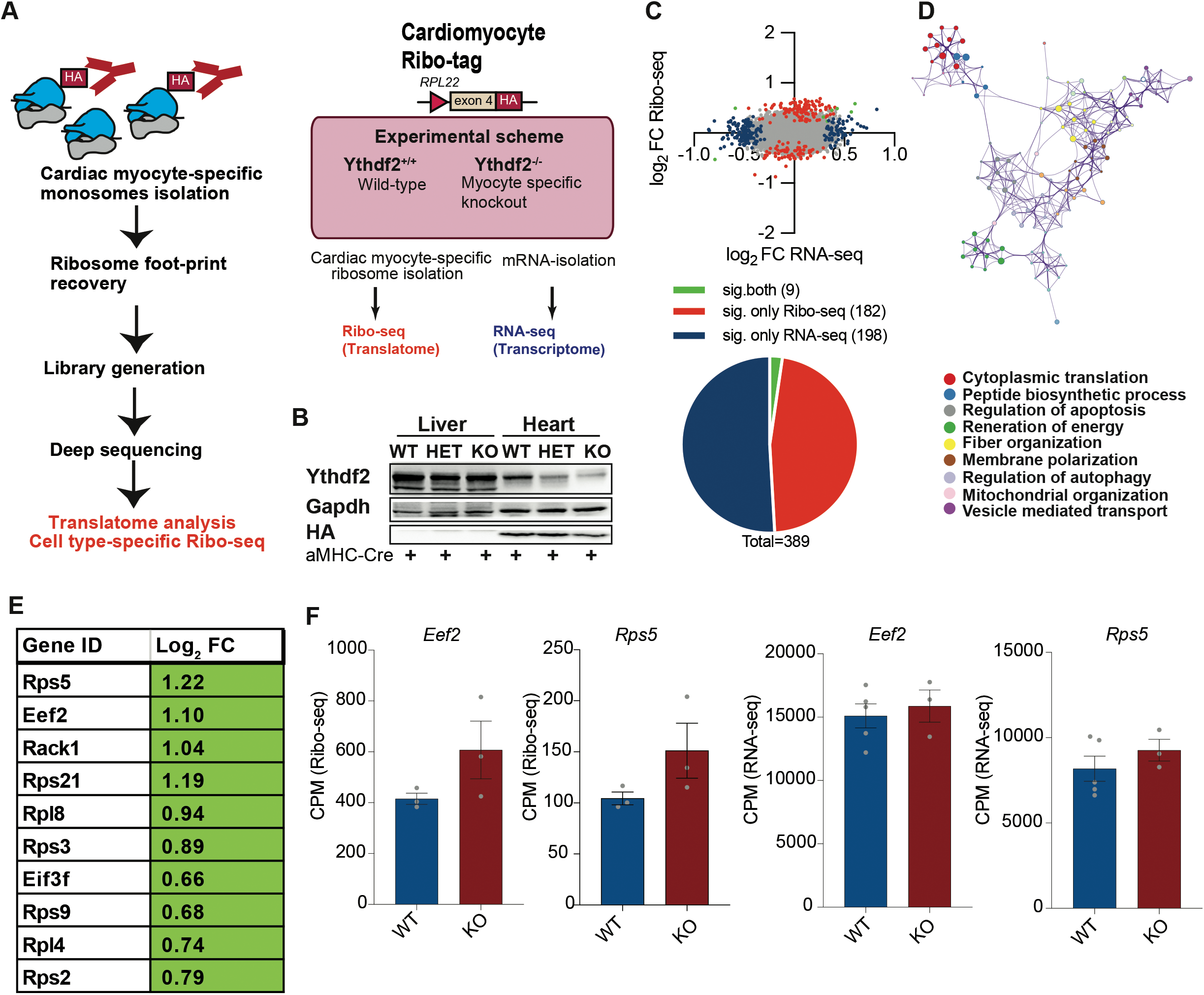
Identification of Ythdf2 targets by Ribo-seq. **A** Schematic of the cell type specific Ribo-seq approach **B** Immunoblot confirming successful Ythdf2 deletion in the heart as well as expression of the HA epitope tag. **C** Scatter blot of log2 fold change of RNA-seq and Ribo-seq (KO/WT), all transcripts are shown in grey, transcriptionally regulated transcripts (blue) with log2 fold change of RNA-seq > 0.5 and p value < 0.05 and translationally regulated transcripts (red) with log2 fold change of Ribo-seq > 0.5 and p value < 0.05. **D** Gene ontology analysis of transcripts that are translationally regulated in KO mouse hearts. **E** Lists of translational regulated genes in Ythdf2 KO mice involved in translational regulation **F** Counts from Ribo-seq and RNA seq for *Eef2* and *RPS5* in KO mice in comparison to WT mice

Parallel, RNA-seq was performed to provide a full view on the transcriptome and the translatome. To identify transcripts that are either regulated on the mRNA or the translational level, log2 fold change of the ratios of normalized counts from Ythdf2 and WT mice were calculated. 83 transcripts were up or downregulated on the mRNA level **(Fig. 4 C, blue)** without parallel changes detected on the translational level by Ribo-seq in Ythdf2 KO vs. WT mice. In contrast, 182 transcripts were differentially translated **(Fig. 4 C, red)** with no significant changes on the mRNA level. Gene ontology (GO) analysis showed that transcripts that are up or downregulated in Ythdf2 KO mice on the translational level are involved in regulating apoptosis, immune system processes, and translation **(Fig. 4 D)**.

The most prominent cluster of enriched transcripts upon loss of Ythdf2 were gene involved in translation and several ribosomal proteins and translational regulators are increased in the Ythdf2 KO mice (**Fig. 4E)**. Since we observed increased translation upon loss of Ythdf2 in cardiomyocytes **(Fig.1)**, we aimed to confirm differential translation of transcripts that are involved in translational regulation. Prominent examples of differentially translated genes upon loss of Ythdf2 were the eucaryotic elongation factor 2 (eEF2) as well as ribosomal proteins such as Rps5. Ribo-seq predicted increased translation of *Eef2* as well as *Rps5* in KO mice compared to WT mice **(Fig. 4 F)** with no changes in mRNA levels. We recently found increased protein levels of Eef2 early after TAC surgery due to elevated translation rates [26]. Consequently, Eef2 was followed up on as an interesting Ythdf2 target involved in cardiac remodeling processes.

To validate the Ribo-seq predictions immunoblots were performed and showed increased protein levels of Eef2in Ythdf2 KO mice compared to WT mice **(Fig. 5 A)**, while mRNA levels were not upregulated **(Fig. 5 B)**.

**Figure 5.**
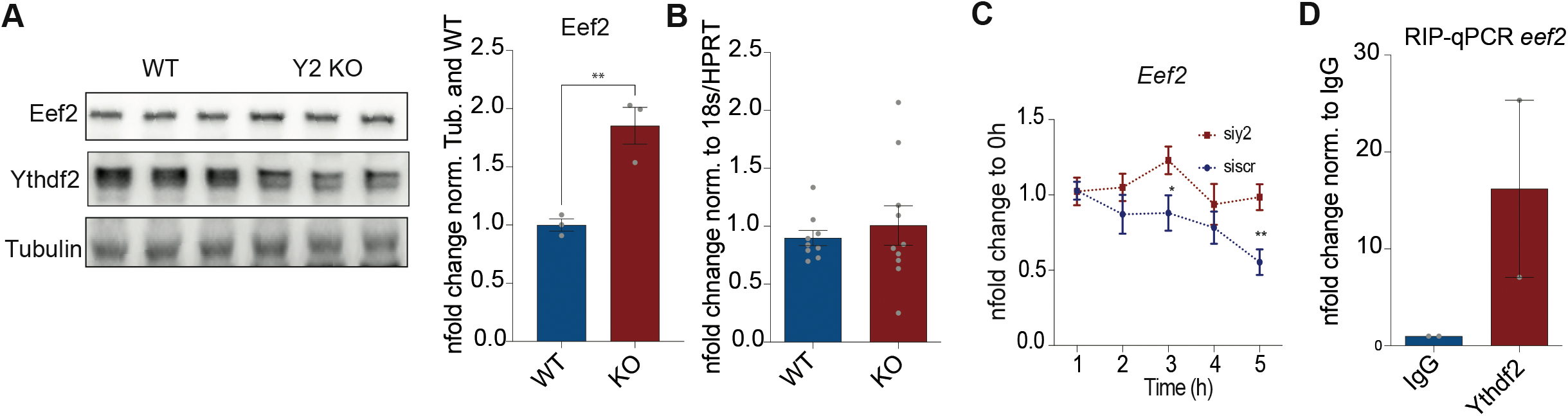
Verification of Ythdf2 dependence of Eef2. **A** Western Blot for Eef2, Ythdf2 and Tubulin and its quantification of WT and KO mice showing the increase of Eef2 protein. **B** *Eef2* mRNA levels are not increased in KO mice in comparison to WT mice, measured by RT-qPCR. **C** *Eef2* transcript stability measured in NRVMs with Ythdf2 knockdown is increased in comparison to scr cells, measured by RT-qPCR+ Actinomycin D Assay. **D** RT-qPCR of RNA from RIPs against Ythdf2 showing *Eef2* mRNA being bound to Ythdf2. Error bar indicates SEM

Ythdf2 was reported to regulate transcript stability by inducing decay. Thus, we hypothesized that Ythdf2 depletion increases Eef2 protein levels by elevating its transcript stability and thereby its availability to translation. We analyzed the effect of Ythdf2 depletion in primary cardiomyocytes on *Eef2* stability using a Actinomycin D assay. *Eef2* mRNA levels decreased in siScr-transfected NRVMs over the treatment time of 6h, while in Ythdf2-depleted cells its levels did not drop to the same extent **(Fig. 5 C)**, suggesting that Ythdf2 indeed regulates *Eef2* transcript stability. Finally, we verified *Eef2* as a Ythdf2 target by immunoprecipitation of Ythdf2-bound RNA followed by qPCR. *Eef2* mRNA was increased in the Ythdf2 RNA precipitation in comparison to the IgG precipitated fraction **(Fig. 5 D)**.

Overall, our study confirmed the importance of m^6^A reader proteins for cardiac function and identified a mechanism how loss of Ythdf2 results in increased translation of *Eef2* which is associated with increased protein synthesis and cellular growth.

## Discussion

Regulation of gene expression in the heart in response to stress is regulated both on the transcriptional and post-transcriptional level [27–29]. Emerging evidence highlights the importance of post-transcriptional regulation of gene expression by non-coding RNAs, RNA binding proteins, and other translational control mechanisms for cardiac function [30, 31]. We and others have shown that direct translational regulation by RNA binding proteins adds an additional level of gene expression to cardiomyocyte homeostasis and pathology [3, 25, 32]. m^6^A is the primary and most abundant base methylation of mRNAs in eukaryotic cells, viruses, and yeast appearing in evolutionary conserved motifs [33]. It was shown in several publications that levels of m^6^A are increased in heart failure, hypertrophic hearts, and isolated cardiomyocytes in response to stress [3–5]. However, very little is known about the m^6^A reader proteins in cardiology. Xu et al. report an increase of Ythdf2 in human heart failure samples and a protective role after overexpression of Ythdf2 in a murine heart failure TAC model [19]. Our study, using a cardiomyocyte specific Ythdf2 KO mouse model, is the first thorough approach unraveling the m^6^A-reader Ythdf2-driven gene expression control in the heart. The loss of Ythdf2 leads to decreased heart function starting at 10 weeks of age in mice and increases gene expression of markers for cardiac remodeling at 30 weeks in comparison to the WT mice. When pressure overload was applied to the left ventricle by TAC surgery, the cardiomyocyte specific loss of Ythdf2 leads to deteriorated heart function, augmented signs of hypertrophy, and worsened fibrosis, indicating a higher loss of cardiomyocytes. Similarly, knockdown of Ythdf2 led to hypertrophic cardiomyocytes. Altogether, our data and previous published reports indicate a protective role of Ythdf2 on cardiac growth and remodeling.

By using parallel Ribo-seq and RNA-seq data from Ythdf2 KO and WT mice, we identified transcripts translationally regulated after loss of Ythdf2. We identified that Eef2, an elongation factor, which is post-transcriptionally upregulated in Ythdf2 KO mice cardiomyocytes and confirmed its increased protein levels without changes on the mRNA level.

Since RIP-qPCR proved that *Eef2* mRNA is bound to Ythdf2 in primary cardiomyocytes and knockdown of Ythdf2 leads to increased *Eef2* mRNA stability, we conclude that Ythdf2 regulates *Eef2* translational activity by binding to an m^6^A site and inducing its decay. The loss of Ythdf2 leads to the extended availability of the *Eef2* transcript for translation. The increased protein level of Eef2 might be an additional driver of cardiomyocyte growth since overall protein synthesis is increased during hypertrophy [22, 34–37]. Obviously, this requires an increase of the translational apparatus like ribosomes, initiation factors, but also elongation factors like Eef2.

In summary, the presentation of the translatome of Ythdf2 KO in combination with the phenotypic analysis of these mice provides new insights into mechanisms of m^6^A-dependent posttranscriptional gene expression control.

## Supplemental Material

**Suppl. Fig. 1 Knockdown of Ythdf2 does not affect the expression of Ythdf1 and 2**

**A** Western Blot for Ythdf1 and **B** Ythdf3 in NRVMs transfected with siRNA against Ythdf2. **C** mRNA expression levels of *Ythdf1* and **D** *Ythdf3* in NRVMs with Ythdf2 KD is not significantly different compared to controls. Error bar indicates SEM

**Suppl. Fig. 2 The cardiomyocyte-specific KO of Ythdf2 increases the expression cardiac remodeling markers in old mice**

**A** Western Blot of lysates from left ventricles (LV) and livers from WT and KO mice with antibodies against Ythdf2, Actin and HA-tag on RPL22. **B-C** mRNA expression levels of **B** *Nppb* are increased while mRNA levels of **C** *Fn1* in left ventricles from 14 weeks old mice WT and Ythdf2 KO mice do not vary significantly, measured by RT-qPCR, * P < 0.05 by unpaired t-test. **D-E** In 30 weeks old Ythdf2 KO mice, the mRNA expression levels of **D** *Nppb* and **E** *Fn1* are significantly increased in comparison to WT mice at the same age, measured by RT-qPCR, * P < 0.05, ** P < 0.001 by unpaired t-test. Error bar indicates SEM **F** Heart weight measurements of WT mice and Ythdf2 KO mice do not vary significantly at about 30 weeks of age.

**Suppl. Fig. 3 The LV/body weight ratio and expression of hypertrophy marker genes are not significantly different in Ythdf2 KO mouse hearts**

**A** Ratios of left ventricle weights (LV) and body weights are increased after TAC surgery in comparison to Sham surgeries, *** P < 0.005, **** P < 0.001 by Kruskal Wallace test. **B-D** mRNA expression levels of the hypertrophic markers **B** *Nppa*, **C** *Nppb* and **D** *Myh7* is increased in KO mice after TAC surgery in comparison to WT mice after TAC surgery, Myh7 measured by RT-qPCR, * P < 0.05, ** P < 0.01, *** P < 0.005, **** P < 0.001 by Kruskal Wallace test. Error bar indicates SEM

**Suppl. Fig. 4 Ribo-Seq of Ythdf2 KO versus WT mice**

**A** Quality control of biological replicates of Ribo-Seq. Segregation of Usable, non-periodic, multimapping and unaligned reads. **B** Scatterplots for pairwise comparisons of the log2 fold change of counts per million mapped reads (CPM) for annotated transcripts (>1 reads in three different Ribo-seq libraries from WT mice). Pearson correlation coefficient r between different libraries > 0.97.

## Methods

### Animals/ Ythdf2 KO mice

All experiments were performed in 10-wk-old male mice unless otherwise indicated. Mice carrying a floxed Ythdf2 allele and the α-MHC Cre Ribo-tag mouse [25] were crossed to obtain a cardiomyocyte specific Ythdf2 KO mouse expressing a HA-tagged RPL22. The mice were housed in a temperature- and humidity-controlled facility with a 12-h light–dark cycle.

At 10 wk of age, male mice underwent TAC (27 gauge needle) or sham operation, as previously described [24]. For echocardiography, the mice were anesthetized with 2% isoflurane and scanned using a Vevo2100 imaging system (Visual Sonics) as previously described [38]. Institutional Animal Care and Use Committee approval was obtained for all animal studies.

### Ribo-seq and RNA libraries

To accurately dissect translation and transcription, both Ribo-seq and RNA-seq libraries were prepared for each biological replicate from aMHC-Cre–positive Ythdf2 KO mice. Cardiomyocyte-specific ribosomes were isolated like previously described[25]. Briefly, the tissue was homogenized in polysome buffer, and the lysate was used as the input for RNA-seq. The remaining lysate was used for direct IP of polysomes. Anti-HA magnetic beads (88836; Thermo Fisher Scientific; 100 μl per heart) were washed with 1,000 μl polysome lysis buffer three times. The lysate was then added to HA magnetic beads and incubated with rotation at 4°C overnight. The beads were then washed three times with 500 μl of high salt buffer (20 mM, Tris pH 7.4, 10 mM MgCl, 300 mM KCl, 2 mM DTT, and 1% Triton X). The washed beads were subjected to RNA extraction for Ribo-seq library construction. Libraries were generated according the to the mammalian ARTseq kit (Illumina). Barcodes were used to perform multiplex sequencing and create sequencing pools containing at least eight different samples and always an equal amount of both RNA and RPF libraries. Sample pools were sequenced on the NextSeq platform using 75-SE sequencing chemistry.

### RiboSeq-Analysis

We followed our previously published protocols (Doroudgar et al.,2019). Adapter removal was done with Flexbar v3.0.3 [39] using standard filtering parameters. Reads were aligned to a custom bowtie2 v2.3.0 [40] ribosomal index were discarded. Remaining reads were then aligned in genome and transcriptome coordinates with a splice-aware aligner (STAR) [41]. We used the EnsEMBL mouse genome assembly GRCm38.p6, where all non-coding regions were excluded, and all fully contained shorter CDSs were collapsed. For Ribo-seq data, only periodic fragment lengths were kept that showed a distinctive triplet periodicity. For the statistical analyses, we use the edgeR package [41]. We only consider data points with read count observations across all replicates. We used an FDR<0.05 and FC=2 (log2 FC=1) as cutoff.

### Primary Cell culture

NRVMs were isolated by enzymatic digestion of 1–3d old neonatal rat hearts and separated from fibroblasts by pre-plating on non-coated dishes before plating. Cardiomyocytes were cultured at the following densities: 6Mio/15cm dish; 3000 000/6 well and 100 000/12 well. Tissue culture plates were coated with 0.1% Gelatin in PBS for 1h. Briefly, cardiac myocytes were plated on gelatin-coated plastic culture wells in DMEM/F-12 medium containing 10% fetal bovine serum. After 24h, the medium was replaced with DMEM/F-12 medium containing 0,5% fetal bovine serum, and the cultures were maintained for 24h before treatment. To mimic pathologic cardiac hypertrophy, cells were treated with 50 μM PE for 24h.

### siRNA induced knockdown of Ythdf2

24h after plating, NRVMs were transfected with either 25 nM scrambled small interfering (si) RNA or siRNA targeting rat Ythdf2 using the transfection agent HiPerfect according to the manufacturer’s protocol. Media containing the transfection solution was replaced by fresh media after 6 h.

### Immunofluorescence staining

NRVMs were grown on gelatin-coated glass chamber slides or for in Cell analysis in 96 black well plate with transparent bottom. Cells were fixed by 4% paraformaldehyde (PFA) for 10 min at room temperature, washed three times with PBS, and permeabilized in PBS with 0.1% Triton-X for 10 min, then blocked in PBS with 10% horse serum for 1h. Primary antibodies diluted in blocking solution (PBS with 10% horse serum) were applied overnight at 4 °C. The next day, cells were washed with PBS and incubated for 1h at RT with secondary antibody (Jackson Laboratories) diluted in blocking solution. After three washing steps in PBS, cells on slides were mounted in Vectashield with 1:10000 DAPI as nuclear staining for slides. NRVMs in 96 black well plates for In Cell Analysis were additionally incubated once with PBS containing 1:1000 DAPI for 5 minutes and then washed again PBS.

### Actinomycin D assay

To inhibit transcription, NRVMs were treated with 2 μM Actinomycin for 1, 2 and 4 and 6 hours. Cells were harvested for RNA isolation and RT-qPCR of Cpeb4 target gene transcripts. Fold changes of ΔCt from Ct of each time point and time point 0 h of Actinomycin D treatment were calculated and non-linear regression curves using GraphPad Prism were modeled in order to determine transcript stability over time.

### Immunoblotting

Samples were combined with the appropriately concentrated form of Laemmli sample buffer and then boiled before SDS-PAGE followed by transfer to PVDF membranes. The membranes were probed with the following antibodies: Ythdf2 (1:1000), Eef2 (1:1000), β-Actin (1:5000), Tubulin (1:5000)

### RNA-Immunoprecipitation

NRVMs were harvested in polysomal lysis buffer (from Invitrogen, 10%Triton, DNase, RNase inhibitor). The lysate was incubated with Ythdf2 AB or IgG as a negative control overnight at 4°C, then for 1h at RT with prepared dynabeads sheep anti rabbit IgG (Invitrogen, #11203D). Coprecipiate was washed once with wash buffer (polysomal buffer, 10% Triton, DNase I), three times with high salt buffer (polysomal buffer + 1M KCl, 10% Triton, DNase I) and once with wash buffer. RNA was eluted by addition of Trizol and following RNA clean-up, followed by cDNA synthesis and RT-qPCR.

### RT-PCR

Quantitative RT-PCR was performed in triplicate measurements on samples from input and RIC eluate RNA and amplified using specific oligonucleotide primers designed to the indicated transcripts primers.

For quantification of *18S* mRNA, *Hprt* and indicated mRNAs reverse transcription followed by RT-qPCR was performed. RT was performed using the iScript™ cDNA Synthesis Kit (Bio-Rad). Amplification was performed in the presence of 1 mM of each oligonucleotide and iTaq Universal SYBR Green Supermix (Bio-Rad). Relative amounts of targets were calculated using the ddCT method. Sequences of oligonucleotides used for quantification of *18S* and *Hprt* mRNA were designed with Primer3 software.

### Statistics

Cell culture experiments were performed at least two times with n = at least two biological replicates (cultures) for each treatment. In vivo experiments were performed on at least three biological replicates (mice) for each treatment. Cell size measurements of NRVMs were performed with at least four biological replicates and repeated twice. Unless otherwise stated, values shown are mean ± SEM. In case of three or more experimental groups, values were tested for normal distribution and then either one-way ANOVA followed by Bonferroni’s post hoc comparisons (normal distribution) or Kruskal Wallis Test followed by Dunn’s multiple comparison was performed. In case of two experimental groups unpaired t-test for normal distributed samples or Mann-Whitney-U-test was performed.

## Data availability

The RNA-seq and Ribo-seq data from this publication have been deposited to the sequence read archive (SRA) database [https://www.ncbi.nlm.nih.gov/sra] and assigned the identifier PRJNA912651.

## Acknowledgments

VK., HAK., NF., and MV. acknowledge the DZHK (German Centre for Cardiovascular Research) Partner Site Heidelberg/Mannheim. MV acknowledges the DFG (German Research Foundation, DFG VO 1659 2/1, DFG VO 1659 2/2, DFG VO 1659 4/1, DFG VO 1659 6/1) and the Boehringer Ingelheim Foundation (Plus 3 Programme). MV and NF acknowledge the CRC 1550.

## Author contributions

VK and JÖ were responsible for collection and/or assembly of data, data analysis and interpretation. VK was responsible for manuscript writing. EV, SH, AM, PG, ZL, VKS, JL, and CS were responsible for collection and/or assembly of data, and data analysis and interpretation. TJ was responsible for RNA-seq and Ribo-seq data analysis and interpretation. HAK, and NF were responsible for data analysis, interpretation, and manuscript writing. MV was responsible for conception and design, data analysis interpretation, manuscript writing and final approval of the manuscript.

