## Supplemental Figures for "Ythdf2 regulates cardiac remodeling through its m^6^A-mRNA target transcripts"

**Supplemental Figure 1**

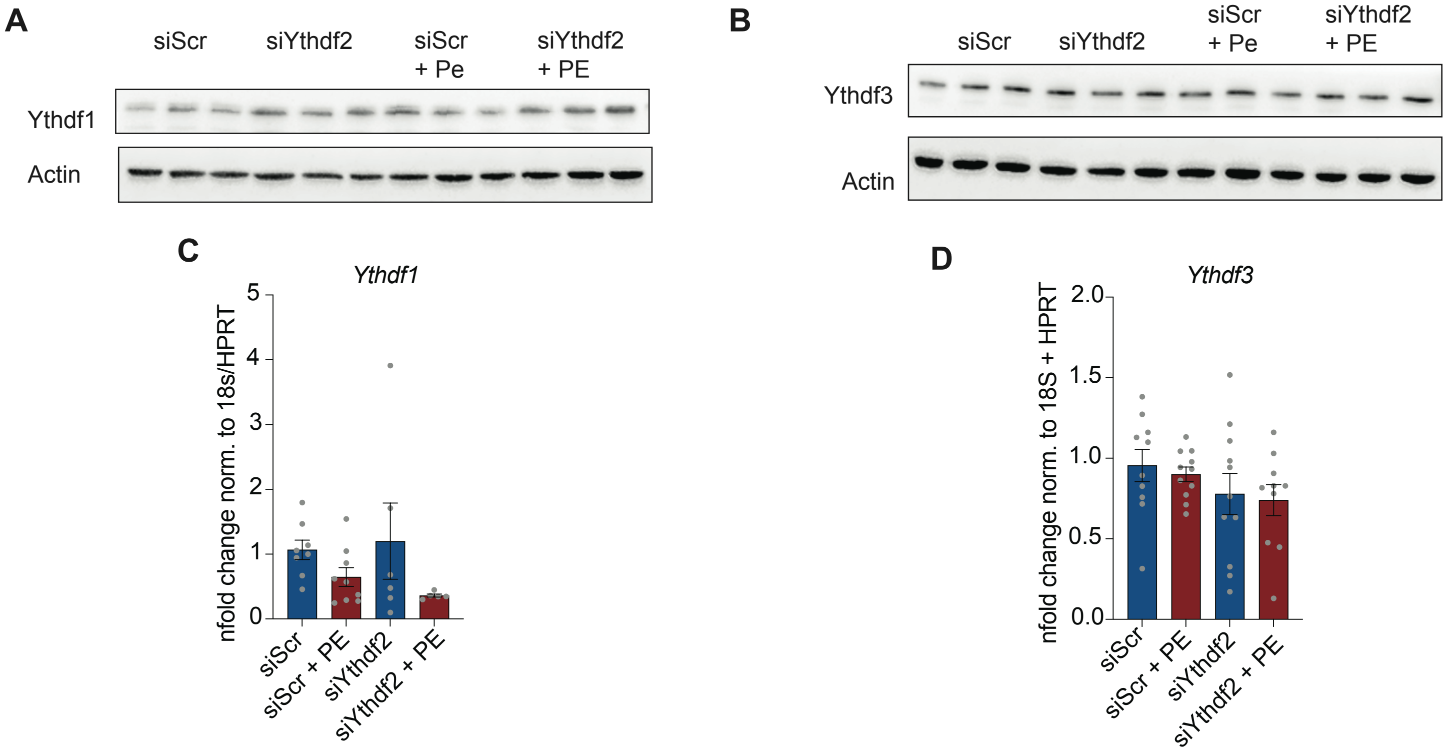

#### Supplemental Figure 2

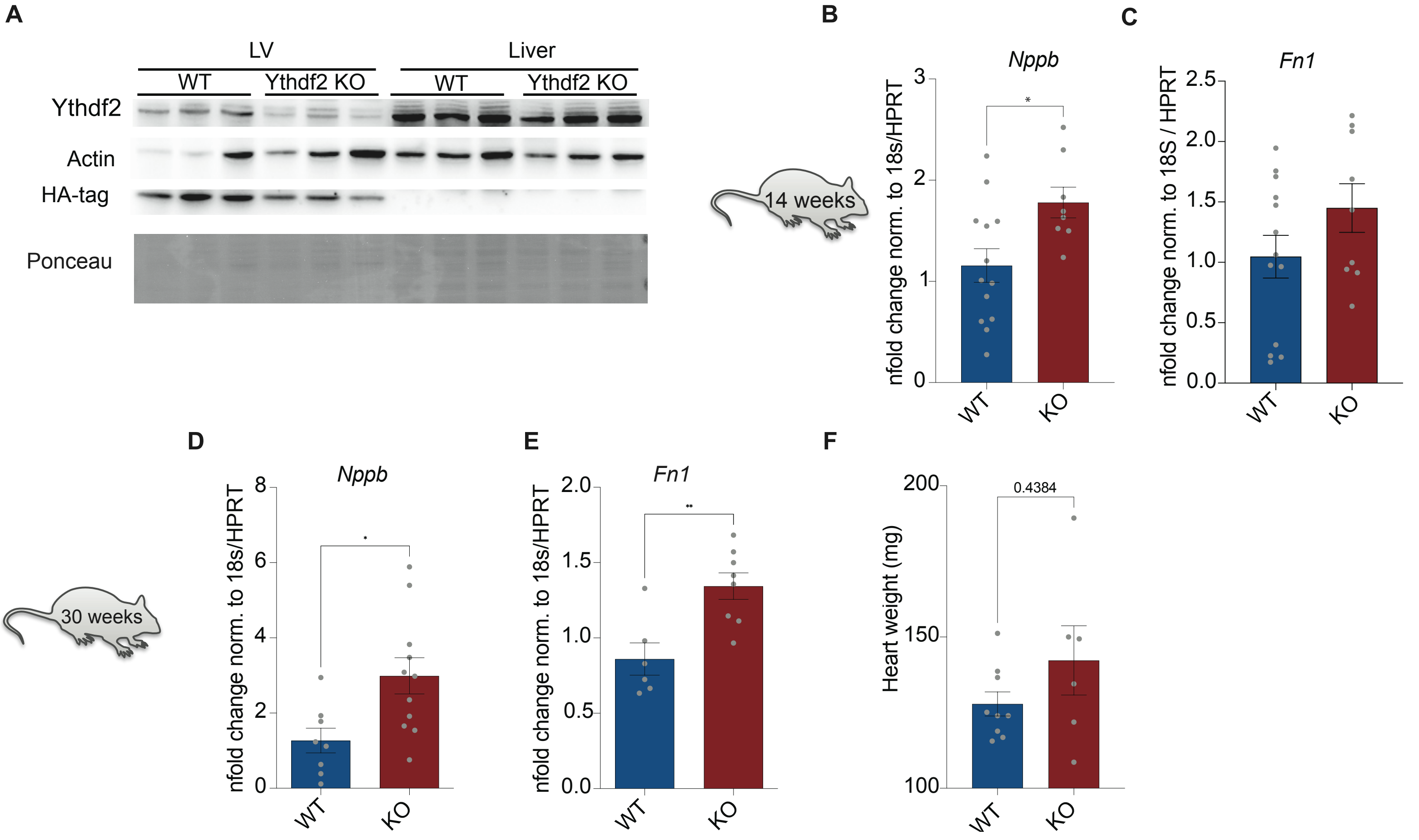

#### Supplemental Figure 3

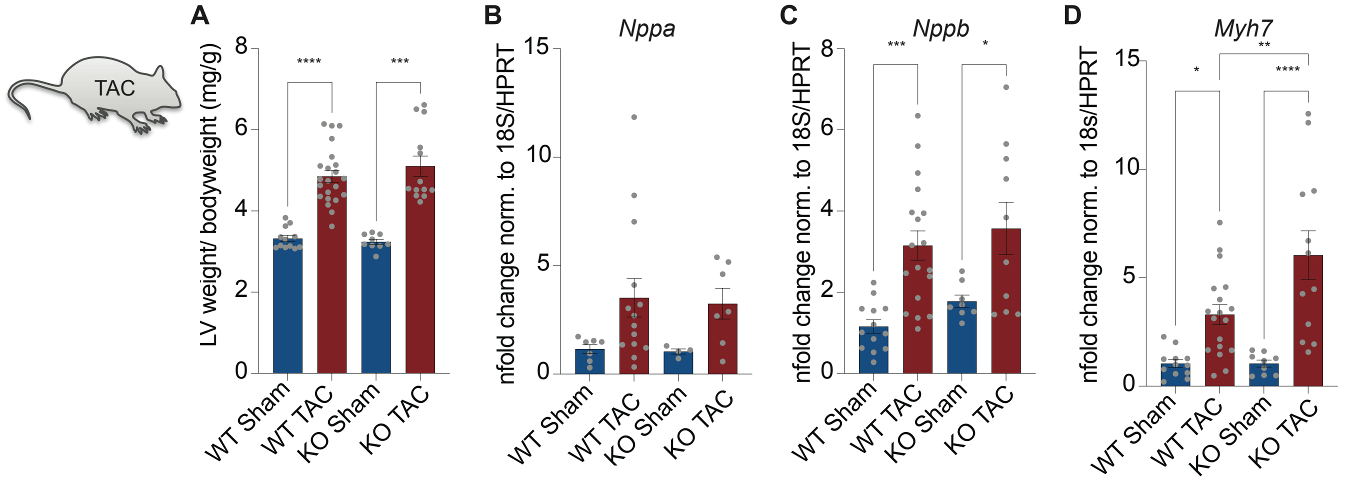

### Supplemental Figure 4

**A** Read filtering counts, no ribosomal matches

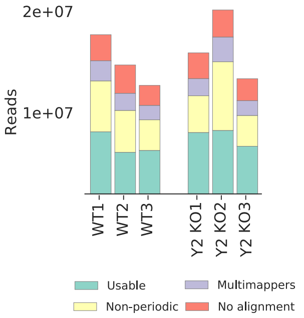


**B**

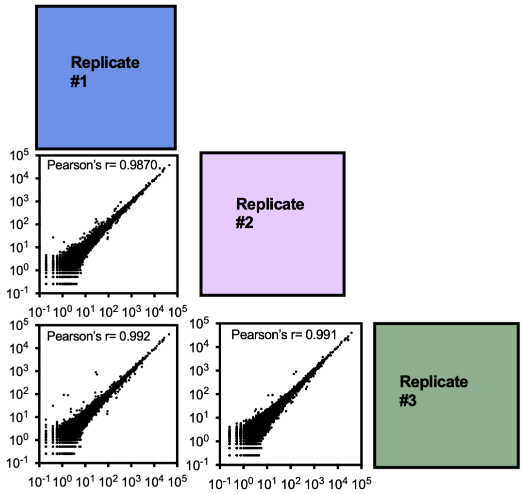
